# CTCF confers local nucleosome resiliency after DNA replication and during mitosis

**DOI:** 10.1101/563619

**Authors:** Nick Owens, Thaleia Papadopoulou, Nicola Festuccia, Alexandra Tachtsidi, Inma Gonzalez, Agnès Dubois, Sandrine Vandoermel-Pournin, Elphège P. Nora, Benoit G. Bruneau, Michel Cohen-Tannoudji, Pablo Navarro

**Affiliations:** Epigenomics, Proliferation and the Identity of Cells, Institut Pasteur, CNRS UMR3738, 25 rue du Docteur Roux, 75015 Paris, France.; Equipe Labellisée LIGUE Contre le Cancer.; Sorbonne Université, Collège Doctoral, F-75005 Paris, France.; Early Mammalian Development and Stem Cell Biology, Institut Pasteur, CNRS UMR 3738, 25 rue du Docteur Roux, 75015 Paris, France.; Gladstone Institutes, San Francisco, CA 94158; Cardiovascular Research institute, University of California, San Francisco, 94158; Department of Pediatrics, University of California, San Francisco, 94143

**Keywords:** nucleosome, transcription factor, mitotic bookmarking, mitosis, replication

## Abstract

The access of Transcription Factors (TFs) to their cognate DNA binding motifs requires a precise control over nucleosome positioning. This is especially important following DNA replication and during mitosis, both resulting in profound changes in nu-cleosome organization over TF binding regions. Using mouse Embryonic Stem (ES) cells, we show that the TF CTCF displaces nucleosomes from its binding site and locally organizes large and phased nucleosomal arrays, not only in interphase steady-state but also immediately after replication and during mitosis. While regions bound by other TFs, such as Oct4 and Sox2, display major rearrangement, the post-replication and mitotic nucleosome organization activity of CTCF is not likely to be unique: Esrrb binding regions are also characterized by persistent nucleosome positioning. Therefore, we propose that selected TFs, such as CTCF and Esrrb, govern the inheritance of nucleosome positioning at gene regulatory regions throughout the ES cell-cycle.

---

Gene regulatory processes are frequently governed by sequence-specific Transcription Factors (TFs) that recognize specific DNA binding motifs **(1)**. TFs are thought to employ different strategies to gain access to DNA **(2–9)**, which in eukaryotes is wrapped around a histone core octamer – the nu-cleosome **(10)**. Whereas nucleosomal DNA is accessible to pioneer TFs **(11, 12)**, TF binding is also associated with the creation of Nucleosome Depleted Regions (NDRs) centered on binding sites and flanked by Nucleosome Ordered Arrays (NOAs) **(13–18)**. However, whether TFs are directly responsible for the establishment and maintenance of these phased nucleosomal structures remains unclear. Hence, understanding the reciprocal relationships between TF binding and nucleosome positioning remains a major goal in the study of gene regulation **(19)**; especially in light of the constraints imposed to chromatin by DNA replication and mitosis. Indeed, both the passage of the replication fork and the mitotic condensation of chromatin are accompanied by a broad eviction of TFs, the loss of NDRs and the disorganization of NOAs, which in turn further impairs TF binding **(20–22)**. In contrast to this general behavior, we show here that selected TFs maintain or rapidly re-establish nucleosome positioning after replication and during mitosis, thereby building nucleosomal resiliency throughout the cell cycle.

The stereotypical NDR/NOA organization at TF binding regions is particularly well illustrated by the genomic binding sites of the zinc finger CCCTC-binding protein (CTCF) **(23, 24)**, a TF involved in chromatin organization and transcriptional control **(25, 26)**. Indeed, we observed a very well defined NOA/NDR/NOA at CTCF binding regions defined by Chromatin Immunoprecipitation followed by sequencing (ChIP-seq) in mouse Embryonic Stem (ES) cells, using Microccocal Nuclease digestion (MNase-seq; Fig. 1A, left), histone H3 ChIP-seq (Fig. 1A, right) and Assay for Transposase-Accessible Chromatin (ATAC-seq; Fig. S1A). This was the case both centering the regions on CTCF motifs (Fig. 1A) and at individual loci (Fig. 1B). CTCF binding footprints, identified as small MNase (<100bp) and ATAC (<150bp) fragments, were also detected within the NDR except upon H3 ChIP (Fig. 1A, C and Fig. S1A). Ranking the regions by their peak height, we observed that the aggregate of CTCF motifs is correlated with CTCF occupancy (Fig. 1C and Fig. S1B). Further, as the motif score and CTCF binding diminishes, the associated footprints decrease and the nucleosome arrays appear less ordered (Fig. 1C). This is particularly well illustrated by the quantitative analysis of the position of the −1 and +1 nucleosomes, which roll inwards and shrink the NDR, concomitantly with increased nucleosome occupancy of the CTCF motif (Fig. 1D). These observations argue for the interaction of CTCF with its cognate binding sites acting as a major force driving the establishment of NDRs/NOAs.

**Fig. 1.**
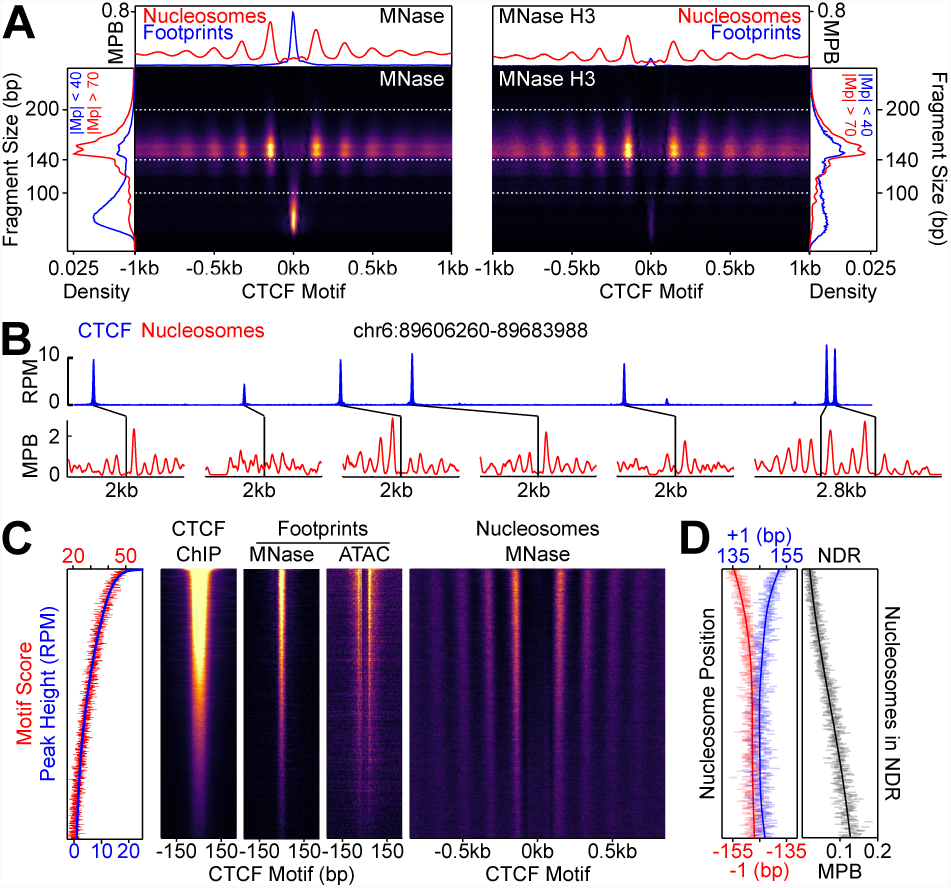
CTCF binding and nucleosome organisation. **(A)** MNase-seq (left) and MNase H3 ChIP-seq (right) V-plots (MNase fragment mid-point vs MNase fragment length) at CTCF binding sites showing +/-1kb surrounding CTCF motifs for fragment sizes in the range 30-250bp. Sidebars indicate densities for fragments with midpoints (Mp) located either within 40bp of the CTCF motif (highlighting CTCF footprints; blue), or at more than 70bp (highlighting nucleosomal fragments; red). Top bar gives metaplots of footprints (fragment length <100bp; blue) and nucleosomes (fragment length within 140-200bp; red). The Y-axis represent fragment midpoints-per-billion (MPB). **(B)** Representative genome snapshot (chr6:89606260-89683988; 78kb) showing CTCF binding in blue (reads per million; RPM) and the associated NDR/NOAs in red (MPB). **(C)** Analysis of CTCF occupancy, CTCF motifs, and nucleosome positioning with decreasing CTCF ChIP-seq peak height: on the left, overlaid of the aggregate of motif scores beneath each CTCF binding region (red) and the height of the corresponding CTCF peaks (blue), analyzed in 100-region bins; the heatmaps, centered to the CTCF motif, correspond to CTCF ChIP-seq signal (the inferred ChIP fragment mid-point is marked), short fragment footprints (marking +/-4bp from midpoint of 1-100bp fragments for MNase-seq and each cut-site of 1-150bp fragments for ATAC-seq shifted inwards by 4bp) and nucleosome-sized fragments (marking the midpoint of 140-200bp fragments). **(D)** Quantitative analysis of the NDR, in 100-site bins descending with CTCF ChIP-seq peak height. Left; the position (in bp) of the median nucleosomal signal (140-200bp MNase-seq fragments with midpoints within +/-70-230 bp from the motif) for the +/-1 nucleosome (blue and red, respectively) per bin and smoothed with Gaussian process regression (GPR; line). Right; mean depth of nucleosomal fragments (MNase H3 ChIP-seq MPB; black) with midpoints in the NDR defined as +/-80bp centered on the motif for each bin and then smoothed with GPR.

To address this functionally, we exploited ES cells expressing CTCF fused to an Auxin-Inducible Degron **(25)** (AID; thereafter CTCF-aid). In accord with the hypomorphic behavior of this line **(25)**, CTCF-aid displayed reduced bind-ing levels (Fig. 2A) that correlated with partially altered nu-cleosome organization compared to wild-type cells (Fig. 2B). Upon short (2h) treatment with the Auxin analogue Indole-3-Acetic Acid (IAA), CTCF-aid expression (Fig. S2A, B) and binding (Fig. 2A) were significantly reduced. This led to a dramatic loss of NOAs, major displacements of the +/-1 nu-cleosomes, and an invasion of the NDR by nucleosomes (Fig. 2B, C), an effect that was observed across different classes of functional genetic features (Fig. 2D). We conclude that CTCF is a major determinant of local nucleosome organiza-tion in steady-state conditions.

**Fig. 2.**
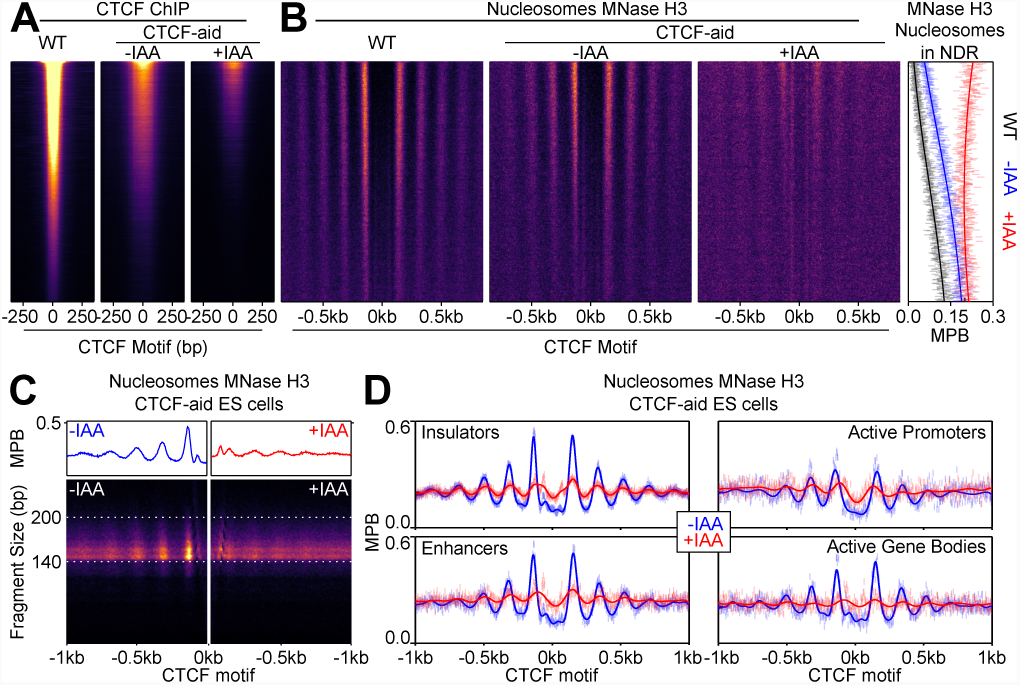
CTCF drives local nucleosome organization. **(A)** CTCF ChIP-seq in wild-type (E14Tg2a; WT) and CTCF-aid −/+ IAA (2h treatment) ES cells, presented as in Fig. 1B and scaled to WT. **(B)** Analysis of nucleosome organization as a function of CTCF binding. Left; heatmaps of nucleosomal fragments of MNase H3 ChIP-seq in WT and CTCF-aid −/+ IAA presented as in Fig. 1B and scaled to WT. Right; nucleosomal signal (MBP) within the NDR (+/-80bp of motif) for WT (black) and CTCF-aid −/+IAA (blue and red, respectively), presented as in Fig. 1B. **(C)** Split V-plot and corresponding metaplot of MNase H3 nucleosomal ChIP-seq signal presented as in Fig. 1A for -IAA (left) and +IAA (right). **(D)** Metaplots of MNase H3 ChIP-seq for -IAA and +IAA centered at CTCF motifs of CTCF peaks intersecting with the indicated ChromHMM categories. Datapoints mark mean MPB per site at each base pair; lines represent Gaussian process regression.

During replication, the chromatin has to be reconstituted downstream of the replication fork **(27, 28)**. While epigenetic marks of both active and inactive chromatin can potentially be propagated from parental to daughter chromatin fibers, only repressed chromatin has been shown to recycle nucleosomes at their previous positions on nascent DNA **(29)**. Hence, at active regions, replication leads to a period during which TFs and nucleosomes enter into direct competition; in Drosophila S2 cells, the reconstitution of specific NDRs/NOAs over active regulatory elements, particularly at enhancers, takes much longer than previously anticipated **(20)**. To address this in ES cells, we used Mapping In vivo Nascent Chromatin with EdU (MINCE-seq; Fig. S3) and generated the average nucleosome profile of several regions (Fig. 3A). We also computed two quantitative parameters: the R2 coefficient revealing the similarity of the nucleosome profiles after replication with the controls (Fig. 3B top panel) and the spectral density assessing nucleosome periodicity (Fig. 3B bottom panel). At ES cell enhancers, centered on p300 summit, we observed an increase in nucleosome density over the NDRs (Fig. 3A) and severely attenuated NOAs (Fig. 3A, B) immediately after replication (2.5 min EdU incorporation; Pulse). In contrast to S2 cells, which even 1h post-replication still display altered nucleosomal structures at enhancers **(20)**, ES cell enhancers are near completely restored during the following hour post-replication (Chase; Fig. 3A, B). More strikingly, CTCF binding regions displayed a remarkable nucleosomal resiliency as only minor changes were appreciable just after replication and, during the following hour, their structure was indistinguishable from the controls (Fig. 3A, B). Given the direct role of CTCF in the control of nucleosome positioning (Fig. 2), and the presence of CTCF on newly synthetized chromatin **(30)**, it is likely that CTCF is capable of rapidly rebinding its sites post-replication to efficiently re-establish NDRs and NOAs. We then explored the impact of replication on nucleosome positioning at regions that we previously showed to be organized around the binding motifs of either Esrrb, or Oct4/Sox2, three master TFs of pluripotency **(22, 31, 32)**. At Esrrb bound regions we observed a prominent NDR centered on the Esrrb motif and two particularly well positioned flanking nucleosomes just after replication. In contrast, at regions bound by the pioneer TFs Oct4/Sox2 **(33)** (as illustrated by the detection of a nucleosome overlapping their motif), nucleosome positioning was profoundly changed upon replication (Fig. 3A, B). Remarkably, for Esrrb we observed that just after replication, the NDR and the positioning of the +/-1 nucle-osomes were more prominent than after 1 hour (Fig. 3A) and the nucleosomes displayed better phasing (Fig. 3B, bottom panel). This suggests that following replication, Esrrb is rapidly rebound at these sites and imposes strong nucleosome positioning, which is subsequently slightly modified by the binding of additional TFs, a phenomenon that we previously described when we compared Esrrb binding in interphase and mitosis **(22)**. In conclusion, while ES cells present fast post-replication nucleosome reorganization compared to S2 cells **(20)**, our analyses indicate that different TFs exhibit drastic differences in their ability to reinstate nucleosome organization (Fig. 3B), with CTCF and Esrrb exhibiting a particularly compelling capacity to restructure nucleosomal arrays within minutes of the passage of the replication fork.

**Fig. 3.**
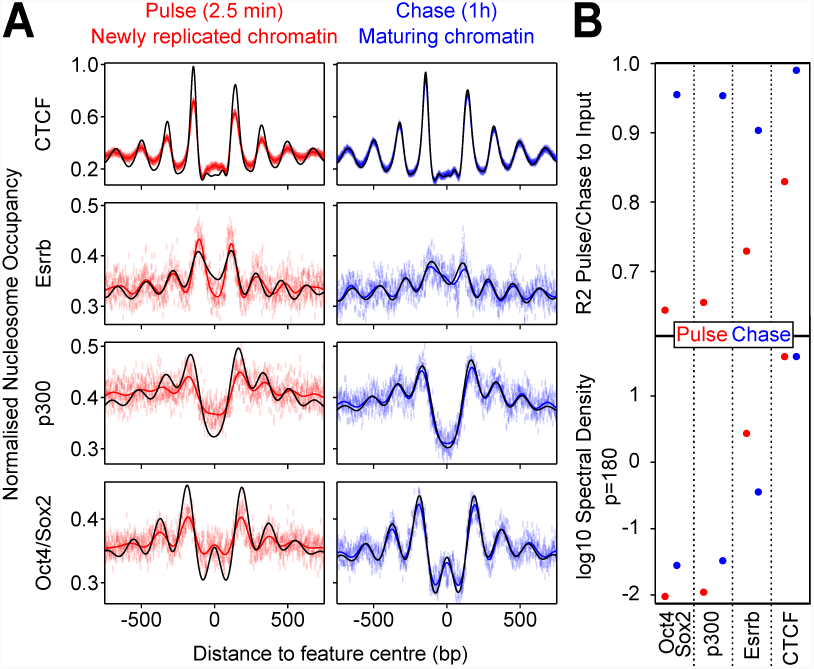
Fast nucleosome positioning at CTCF and Esrrb binding regions after replication. **(A)** Metaplots of nucleosomal fragments at CTCF (motif centered), Esrrb (motif centered), enhancer (p300 summit centered) and Oct4/Sox2 (motif centered) sites in MPB normalized to input/steady-state control for MINCE-seq pulse (replicating chromatin; 2.5min) and chase (maturing chromatin; 1h). Datapoints represent mean MPB per site at each base pair and lines show Gaussian process regression. Black lines display the relevant input/steady-state control. **(B)** Quantification of reconstitution of steady-state nucleosomal order. Top: R^2^ between relevant pulse/chase and controls. Bottom: log10 spectral density at p=180 (nucleosome + linker) of Gaussian process squared exponential covariance function, with optimized hyperparameters.

Mitosis is accompanied by the condensation of the chromatin and the eviction of many TFs from their DNA targets **(34, 35)**. This mitotic absence of TF binding correlates with a drastic reorganization of the nucleosomal landscape at regulatory elements **(22)**, including CTCF-bound regions as assessed in human somatic cells **(21)**. However, in mitotic ES cells we observed the typical NDR/NOA structure at CTCF binding regions (Fig. S4A). Therefore, we hypothesized that CTCF acts as a mitotic bookmarking factor, capable of sitespecific DNA binding in mitotic ES cells as we showed previously for Esrrb **(22, 36)**. In agreement, CTCF ChIP-seq established that this TF binds over half of its interphase targets in mitotic cells (Fig. 4A, B, and Fig. S4B). Cohesin, a recurrent partner of CTCF in interphase **(26)**, was found fully evicted from its targets in mitosis **(37, 38)** (Fig. 4B and Fig. S4B, C), underscoring the specificity of our observation for CTCF. Notably, we observed that nearly all regions displaying robust enrichment for Cohesin in interphase are bookmarked by CTCF, even though Cohesin accumulation is not a prerequisite for CTCF bookmarking (Fig. 4B, C and Fig. S4B, D). Moreover, we found the existence of high quality CTCF motifs to be a better indicator of binding in mitosis than in interphase (Fig. S4D). This suggests that the conditions for CTCF binding are more stringent in mitosis than in interphase. As we showed in interphase (Fig. 1), mitotic CTCF binding was also directly associated with nucleosome organization as assessed both globally (the progressive reduc-tion of mitotic CTCF binding correlates with a gradual loss of NDRs and NOAs; Fig. 4A), and at individual loci (Fig. 4B). Accordingly, when CTCF binding regions were split as bookmarked or lost in mitosis, we could confirm that the nucleosomes are less well positioned (Fig. 4C), phased (Fig. 4D), and exhibit clear displacements toward the motif (Fig. 4D, E), upon the mitotic eviction of CTCF. Overall, this indicates that CTCF is a mitotic bookmarking TF that preserves nucleosome organization during mitosis. Nevertheless, even CTCF bookmarked regions presented a strong displacement inwards of the +1 nucleosome (22bp for +1 versus 3bp for −1 nucleosome; Fig. 4A) and a more moderate shift of all following nucleosomes (Fig. 4D, E). This indicates that the constraints imposed on the nucleosomes by CTCF are slightly different in interphase and in mitosis. Since in interphase CTCF sites that do not bind Cohesin do not show this repositioning of the +1 nucleosome (Fig. S4E), it cannot be explained by the mitotic loss of Cohesin. Additional factors therefore influence nucleosome organization at CTCF binding regions in either interphase or mitosis. Next, we aimed at exploiting CTCF-aid ES cells to more directly address the impact of a loss of mitotic CTCF bookmarking. In mitosis, we observed that the hypomorphic character of CTCF-aid binding was amplified; CTCF-aid was barely detectable at regions binding CTCF in wild-type mitotic cells (Fig. S4G). This global loss of mitotic bookmarking was associated with the acquisition of nucleosomal properties characteristic of regions losing mitotic CTCF in wild-type cells: nucleosome positioning was strongly attenuated (Fig. 4D); nucleosomes, especially upstream of the motif, shifted inwards (55bp upstream versus 21bp downstream; Fig. 4D, E); the NDRs were partially invaded by nucleosomes (Fig. S4H). These changes in nucleosome organization were only minimally increased upon IAA treatment, which leads to further invasion of the NDR by nucleosomes (Fig. S4H). We conclude, therefore, that CTCF behaves as a canonical bookmarking factor in ES cells and actively maintains nucleosome organization, mirroring our previous data on Esrrb **(22)**. While this finding contradicts recent data in human somatic cell lines **(21)**, our analyses in mouse somatic cell lines (NIH3T3 and C2C12) revealed no or limited evidence of mitotic bookmarking activity by CTCF (Fig. S4I). Therefore, even though CTCF can decorate mitotic chromosomes globally, as revealed by microscopy in cell lines and embryos (Fig. S5), this is not necessarily translated into mitotic bookmarking capacity in every cell type. This cell type-specific bookmarking activity of CTCF reconciles our and previous reports **(21, 39, 40)** and further indicates that the mitotic bookmarking activity of CTCF is a developmentally regulated phenomenon. Whether CTCF bookmarking is strictly specific to ES cells, when during development does it lose this activity, and how this impacts long-range chromatin interactions in mitosis and early in the following interphase, represent clear lines of investigation for the future.

**Fig. 4.**
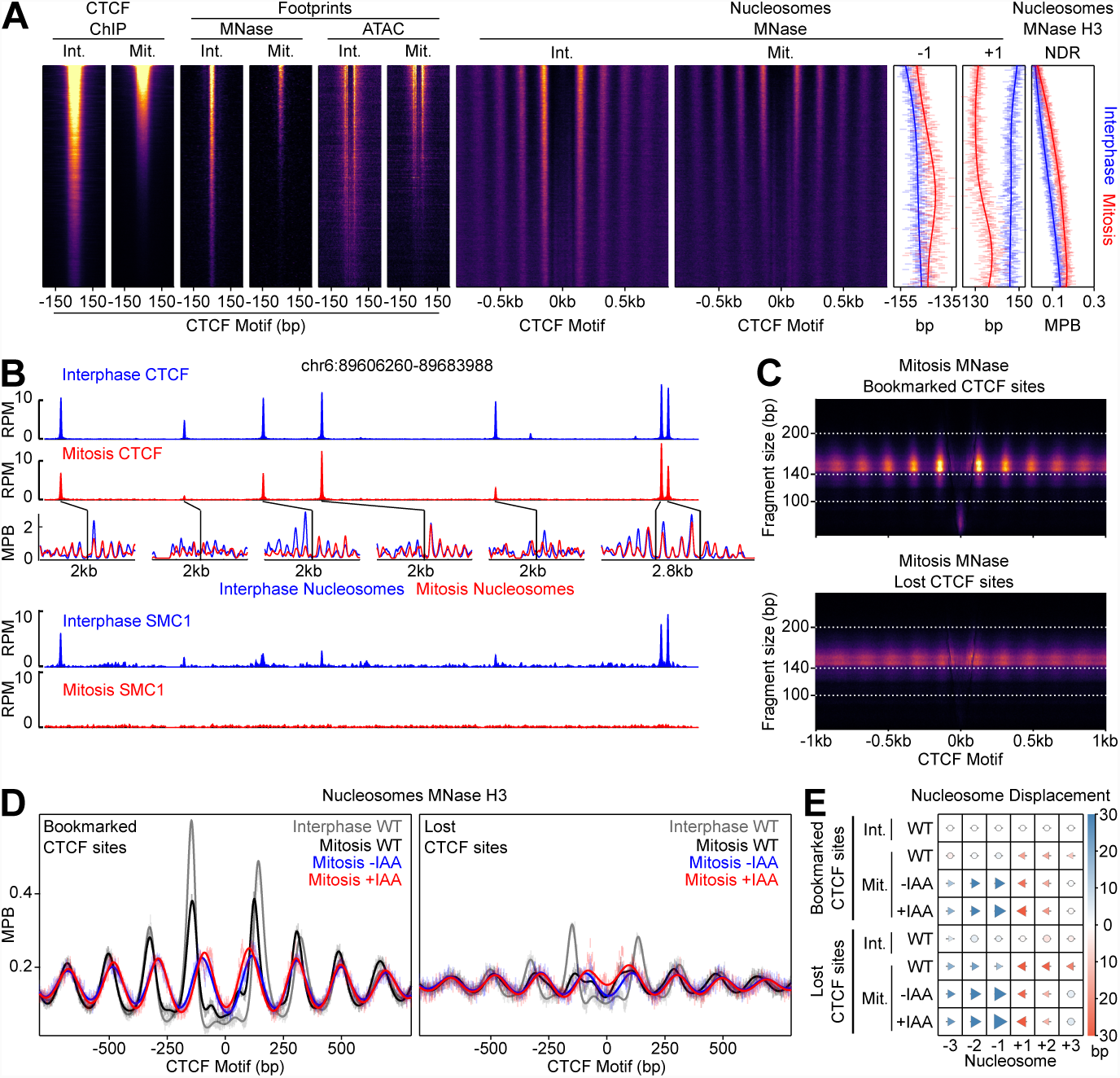
CTCF is a mitotic bookmarking factor in ES cells. **(A)** Interphase (Int.) and mitosis (Mit.) data presented as in Fig. 1C for CTCF ChIP-seq; MNase-seq and ATAC-seq footprints; MNase-seq nucleosomal signal; +/-1 nucleosome positions and NDR signal (MNase H3 ChIP-seq). The regions are ordered by descending mitotic ChIP-seq peak height. All heatmaps are scaled to interphase. **(B)** Representative genome snapshot (chr6:89606260-89683988; 78kb; as in Fig. 1B) for CTCF ChIP-seq (RPM), MNase-seq nucleosome fragments (MPB) and Cohesin (SMC1) ChIP-seq (RPM) in interphase (blue) and mitosis (red). **(C)** Mitosis MNase V-plots for CTCF Bookmarked (top) and Lost regions (bottom), presented as in Fig. 1A. **(D)** MNase H3 ChIP-seq metaplots at CTCF bookmarked (left) and lost sites (right) centered on CTCF motifs for wild-type interphase and mitosis (grey and black, respectively) and for mitosis CTCF-aid −/+ IAA (blue and red, respectively). **(E)** Mean +/-1,2,3 nucleosome positions relative to wild-type interphase, for datasets shown in (D). Circles denote nucleosome movements < 5bp, arrow direction, size and color describes movements > 5 bp.

Altogether, this work reveals that CTCF precisely positions the nucleosomes at steady-state in interphase and, more strikingly, during the challenges posed to the chromatin by replication and mitosis. It is particularly remarkable, moreover, that both CTCF and Esrrb binding regions containing high quality motifs display similar properties during replication and mitosis, whereas regions bound by Oct4/Sox2 do not **(22)**. Whether every mitotic bookmarking TF positions nucleosomes not only in mitosis but also immediately after replication, and whether in doing so they facilitate reassembly of regulatory complexes in nascent chromatin and in daughter cells **(41)**, needs now to be addressed. Indeed, a comprehensive investigation of TFs building local nucleosome resiliency throughout the cell cycle will identify their role in proliferative and developmental processes. This will ultimately illuminate whether their activity bypasses the requirement for a robust epigenetic memory of active regulatory elements **(42)**, particularly in cell types with increased plasticity such as ES cells **(35, 43)**.

## Acknowledgements.

The authors acknowledge the Imagopole France–BioImaging infrastructure, supported by the French National Research Agency (ANR 10-INSB-04-01, Investments for the Future), for advice and access to the UltraVIEW VOX system. We also acknowledge the Transcriptome and EpiGenome, BioMics, Center for Innovation and Technological Research of the Institut Pasteur for NGS. We thank Steven Henikoff and Srinivas Ramachandran for advice on setting up MINCE-seq; Andrea Voigt for sharing protocols for EdU incorporation and click reactions; Marlies Oomen and Job Dekker for discussions and critical reading of the manuscript. This work was supported by recurrent funding from the Institut Pasteur, the CNRS, and Revive (Investissement d’Avenir; ANR-10-LABX-73). E.P.N. was supported by EMBO (ALTF523-2013), HSFP, and the Roddenberry Stem Cell Center at Gladstone. N.O. is supported by Revive. P.N. acknowledges financial support from the Fondation Schlumberger (FRM FSER 2017), the Agence Nationale de la Recherche (ANR 16 CE12 0004 01 MITMAT), the Ligue contre le Cancer (LNCC EL2018 NAVARRO) and the European Research Council (ERC-CoG-2017 BIND).

## Author contributions

The large majority of experiments were performed by T.P. and analyzed by N.O. Experimental help was provided by N.F. and I.G. (mitotic ChIP-seq), A.T. (live imaging) and A.D. (cell culture). Embryo work was performed by S.V.P. and M.C.T. CTCF-aid ES cells were generated by E.P.N. and B.G.B. The project was conceived and supervised by P.N., who wrote the manuscript together with N.O. and T.P.

## Declaration of interests

The authors declare no competing interests.

## SUPPLEMENTARY INFORMATION

Five Supplementary Figures and Methods accompany this manuscript online:

**Fig.S1:** Additional information on CTCF binding in interphase.

**Fig.S2:** Auxin-induced degradation of CTCF-aid.

**Fig.S3:** MINCE-seq controls.

**Fig.S4:** CTCF and SMC1 binding in interphase and mitosis.

**Fig.S5:** Global behavior of CTCF in mitotic cells.

