## Supplementary Figures for "CTCF confers local nucleosome resiliency after DNA replication and during mitosis"

This file contains 5 Supplementary Figures

Supplementary Tables and Methods are available online

**Figure S1:** Additional information on CTCF binding in interphase.

**Figure S2:** Auxin-induced degradation of CTCF-aid in interphase and mitosis.

**Figure S3:** MINCE-seq controls.

**Figure S4:** CTCF and SMC1 binding in interphase and mitosis.

**Figure S5:** Global behavior of CTCF in mitotic cells.

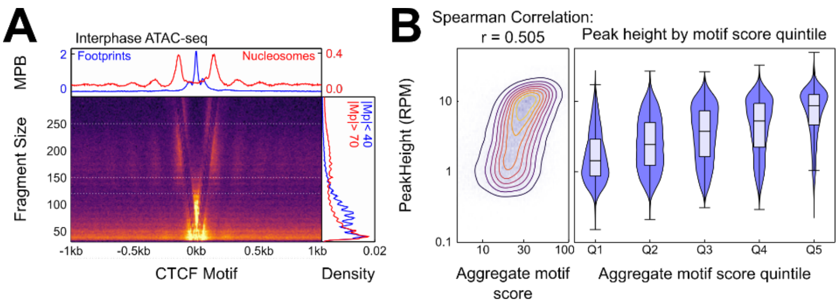


**Figure S1: Additional information on CTCF binding in interphase. (A)** V-plot of ATAC-seq data presented as in Fig. 1A, except that color scale is square root normalized and it covers fragments in the range [30, 300] bp. Footprints calculated from fragment < 150bp, and nucleosome signal from fragments in [150, 250] bp. **(B)** Relationship between CTCF aggregate motif score (the sum of all FIMO motif scores within a peak) and CTCF interphase peak height in (RPM), displayed as contour/heatmap (left) and violin plot of peak height for the quintiles of motif score.

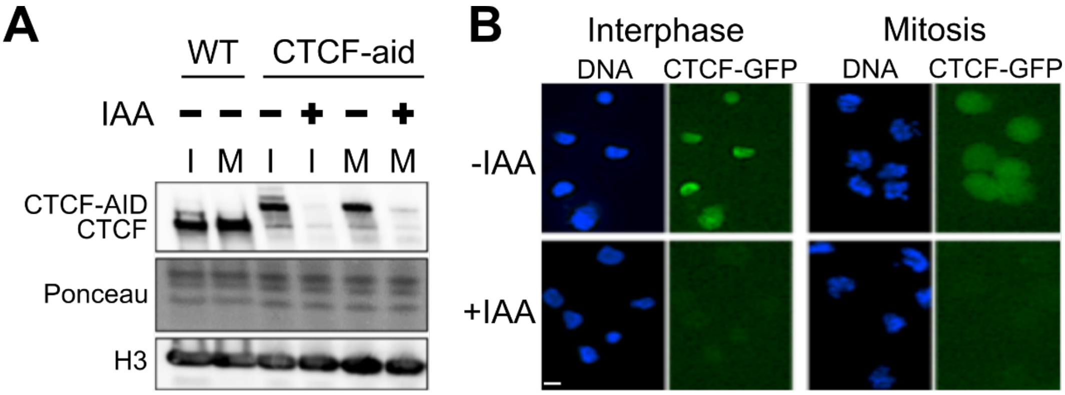


**Figure S2: Auxin-induced degradation of CTCF-aid in interphase and mitosis.** Expression of CTCF-aid, which is fused in frame to GFP (25), by Western Blot **(A)** and Imaging **(B)** after 2h of IAA treatment. Both during interphase (I) and in mitotic (M) cells, CTCF-aid was efficiently degraded.

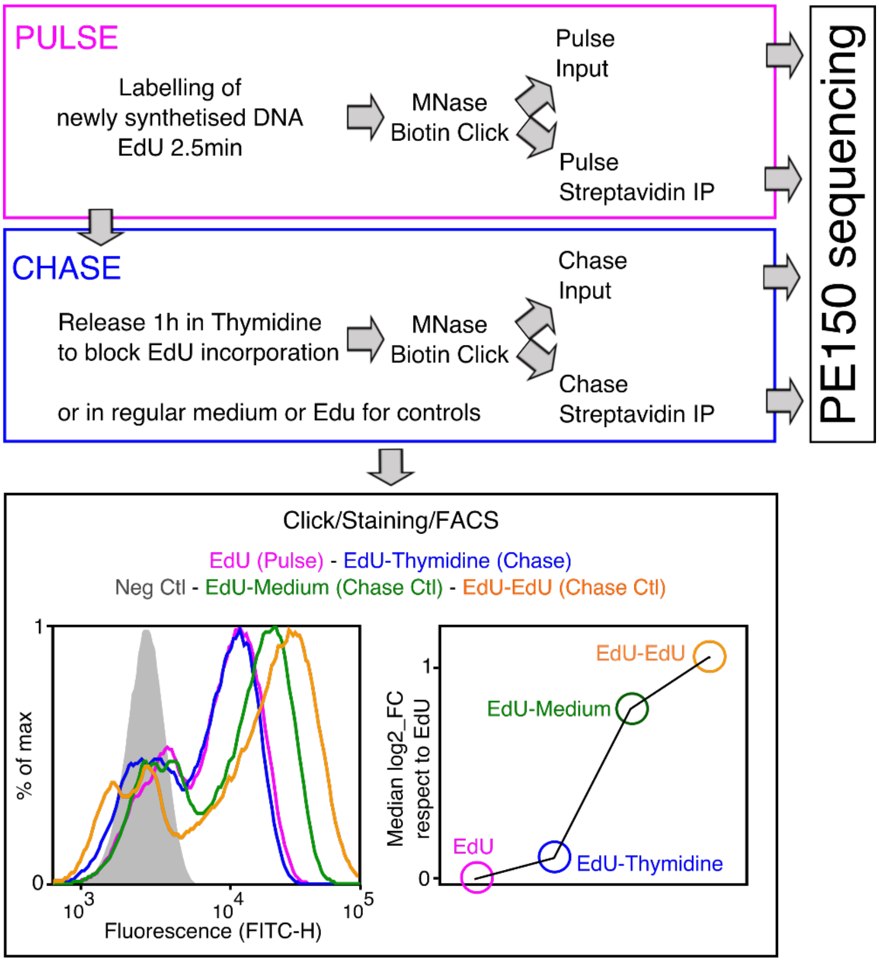


**Figure S3: MINCE-seq controls.** Depiction of the main steps of our protocol, adapted from previous work (20, 44, 45). In ES cells we could use extremely short pulses (2.5min) leading to around 60% of cells labelled by EdU as shown by FACS. After the pulse, the cells were washed and released in Thymidine to block the incorporation of remnant EdU. These samples were processed as indicated to generate sequencing libraries. The efficiency of the chase regarding the lack of further EdU incorporation was assessed by releasing EdU-treated cells in either regular medium or in EdU-containing medium, which lead to readily detectable increases in EdU incorporation, as shown by FACS.

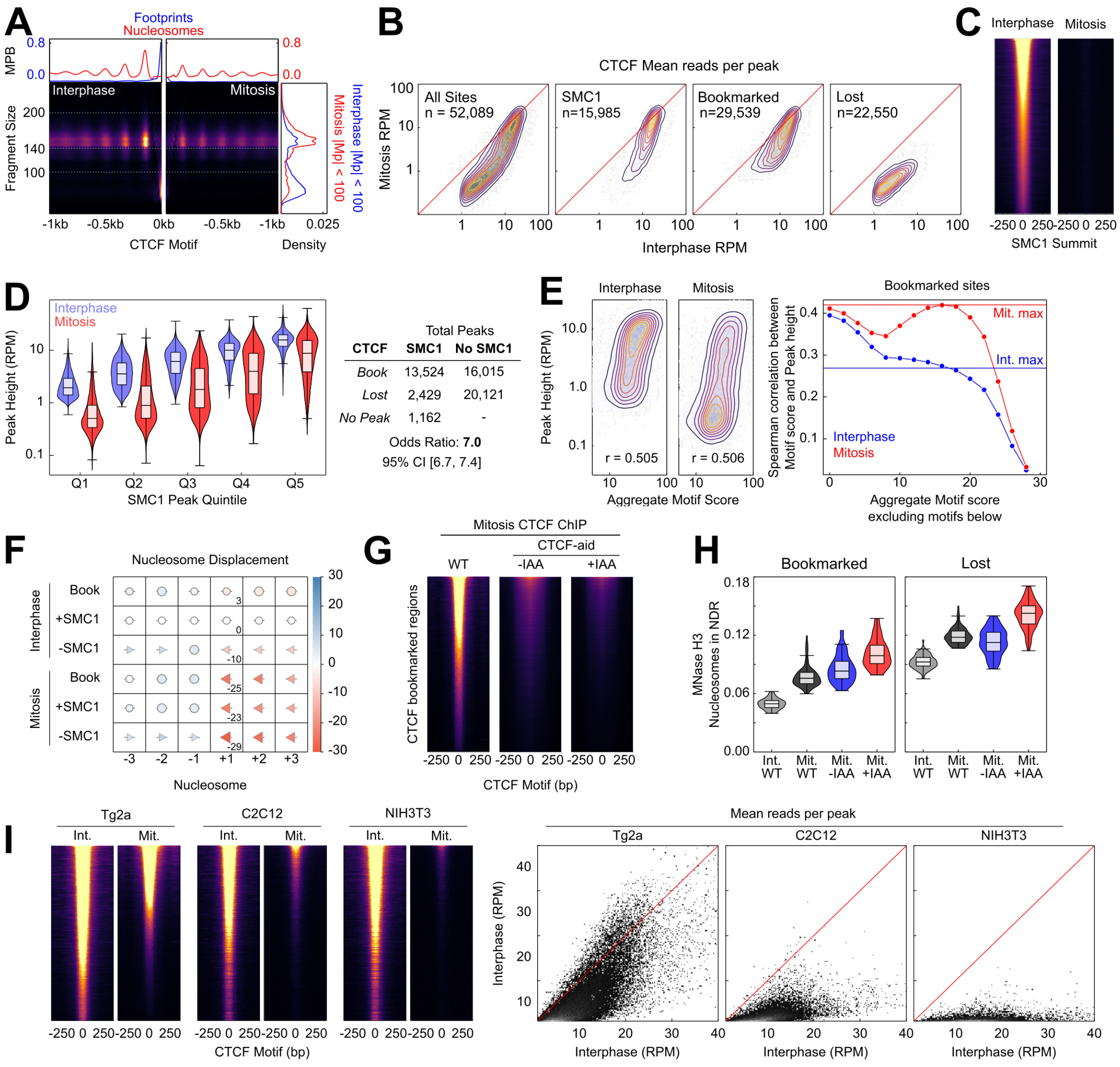


**Figure S4: CTCF and SMC1 binding in interphase and mitosis. (A)** Split V-plot of MNase data at CTCF binding sites centered on maximal CTCF motif in interphase (left) and mitosis (right) for fragments with midpoints [-1kb, 0kb] of CTCF motif (as Fig. 2C). **(B)** Heatmap/contour plots of mean reads per CTCF peak in interphase and mitosis for all peaks, SMC1 intersecting peaks, bookmarked peaks and lost peaks as indicated. **(C)** SMC1 ChIP-seq in interphase and mitosis at SMC1 interphase peaks centered on SMC1 summit, scaled to interphase signal. **(D)** Relationship between SMC1 and CTCF occupancy; left: violin plots of CTCF peak height (RPM) in interphase and mitosis for SMC1 peak height quintiles; right: contingency table of CTCF peaks with and without SMC1 against those bookmarked and lost. Fisher p-value < 2.2e-16. **(E)** Relationship between CTCF aggregate motif score (the sum of all FIMO motif scores within a peak) and CTCF peak heights in interphase and mitosis. Left: contour/heatmap as Fig. S1B. Right: Spearman correlation coefficient between interphase and mitosis peak height and a thresholded aggregate FIMO score in which all motifs less than or equal to the value on the horizontal axis were excluded. Horizontal lines indicate the Spearman correlation between the maximum scoring CTCF motif within each peak and peak height. Mitosis peak heights have a greater dependence on high scoring motifs. Interphase peak heights share the greatest correlation with the sum of all motifs, regardless of FIMO score. **(F)** Nucleosome displacement (as Fig. 4E) of bookmarked, SMC1 positive and SMC1 negative peaks in interphase and mitosis relative to SMC1 positive interphase peaks. **(G)** CTCF ChIP-seq in mitosis for wildtype (WT), CTCF-aid -IAA and +IAA, at CTCF bookmarked sites. All heatmaps scaled to WT interphase. Due to low signal CTCF-aid ChIP-seq marks entire inferred fragment rather than midpoint. **(H)** Nucleosome signal in NDR at bookmarked (left) and lost sites (right), for datasets shown in Fig. 4D; violins and boxplots indicate the variation in signal over each base pair in [-40, 40]bp of the motif. **(I)** CTCF ChIP-seq in interphase and mitosis in E14Tg2a (Tg2a), C2C12 and NIH3T3 at Tg2a peaks; heatmaps (left) mark entire inferred ChIP-seq fragment and are scaled to Tg2a; scatterplots (right) show linear scale reads per million per peak in interphase and mitosis.

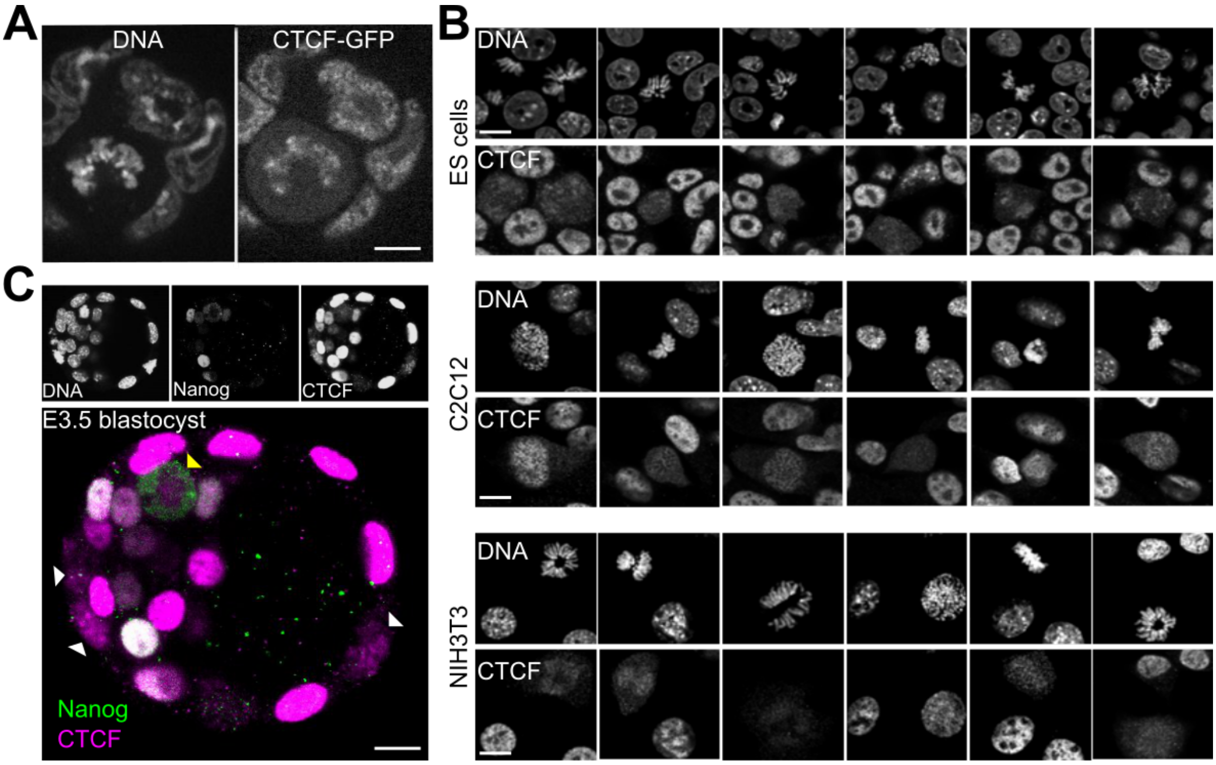


**Figure S5: Global behavior of CTCF in mitotic cells. (A)** Live imaging of CTCF-aid ES cells that also carry a GFP in frame with CTCF-aid (25); coating of the mitotic chromosomes was readily detectable. **(B)** In contrast to other TFs that require alternative fixation approaches based on DSG or Glyoxal to capture the bulk of interactions with the mitotic chromosomes (22), CTCF could be crosslinked with formaldehyde and no major differences were observed with DSG or Glyoxal (not shown). In the three cell lines that we tested (ES cells, C2C12 and NIH3T3) we observed cell-to-cell heterogeneity in respect to CTCF retention during mitosis. The signal observed in C2C12 and NIH3T3 tends to be less prominent than in ES cells and more variable, with some cells exhibiting a clear chromosomal exclusion. **(C)** Out of 7 embryos (of which 5 were treated with nocodazole), we could observe 46 mitoses: 40 in the trophectoderm (TE) and 6 in the inner cell mass. In the TE, 37 mitoses displayed clear coating of the chromosomes by CTCF and the other 3, dimmer signal; the signal observed in 6 mitotic cells of the ICM was more diffused although not excluded from the chromosomes as seen for Nanog.
