## Supplementary material for "CTCF confers local nucleosome resiliency after DNA replication and during mitosis": Methods

### Experimental Methods

#### Cell Culture and mitotic preparations

ES cells (wild-type E14Tg2a and CTCF-aid derivatives) were cultured on 0.1% gelatine (SIGMA, G1890-100G) in DMEM+GlutaMax-I (Gibco, 31966-021), 10% FCS (Gibco 10270-098), 100µM β-mercaptoethanol (Gibco, 31350-010), 1×MEM non-essential amino acids (Gibco, 1140-035) and 10ng/ml recombinant LIF (MILTENYI BIOTEC, 130-099-895). ES cells were passaged 1:10 every 2–3 days. NIH3T3 and C2C12 cells were cultured in DMEM + GlutaMax-I supplemented with 10% FCS. NIH3T3 and C2C12 were passaged 1:15 and 1/20, respectively, every 3 days. To obtain mitotic ES cells (>95% purity as assessed by DAPI staining and microscopy), we used a nocodazole shake-off approach, as described before (*1*). To deplete CTCF-aid in interphase, cells were treated with 0.5mM auxin (IAA Sigma, I5148) for 2h; to deplete CTCF-aid in mitosis, the cells were first treated with nocodazole for 4h and then with nocodazole and IAA for 2h, after which they were harvested by shake-off. C2C12 cells were synchronised like ES cells except that slightly longer nocodazole treatment (7h) was required to improve the yield. NIH3T3 cells were synchronised with a triple approach. First, $1.5x10^6$ cells were seeded in 150mm dishes and grown with 2mM thymidine for 18h to enrich in cells arrested at the G1/S transition. After two washes with PBS, they were released for 8h in regular medium before being arrested again for 17h with 2mM Thymidine. After the double thymidine block the cells were washed twice with PBS and released in regular medium for 7h. Finally, cells were incubated with nocodazole for 6h and mitotic cells isolated by gentle tapping of the dishes against an immobilised surface.

**MINCE-seq**

The protocol was developed based on three existing protocols (*2–4*); for the CLICK reaction we used radical-free aerobic conditions, as previously described (*5*).

*Pulse/Chase with EdU*: $15x10^6$ live cells were seeded per 150mm dish 24h prior to the beginning of each experiment (3 dishes per condition). The cells were then incubated with pre-warmed medium containing 5μM EdU for 2.5min at 37$^o$C. Next, the cells were either harvested and fixed (pulse) or, in parallel, washed with pre-warmed medium and incubated for 1h in the presence of 100μM Thymidine for 1h at 37$^o$C (chase).

These samples and the respective controls (Fig. S3) were checked by FACS after fixation (4% para-formaldehyde, 15min at room temperature protected from light), permeabilization (PBS 1% BSA/ 0.1% Saponin for 15min at room temperature), incubation with 100μl CLICK reaction Mix (see below), and labelling with Streptavidin-Alexa Fluor™ 488 (S32354, Thermo Fisher; diluted 1:1300 in permeabilisation solution) for 1h at room temperature in the dark.

*Microccocal Nuclease digestion (MNase)*: after trypsinisation, $120\times10^6$ cells per sample were fixed with 1% formaldehyde (Thermo, 28908; 10min at room temperature with occasional mixing) and quenched with 0.125M Glycine (5min). Fixed cells, after being washed twice with PBS, were lysed for 10min on a rotating wheel at 4$^o$ C in 5mL of lysis buffer (50mM HEPES pH 7.8/ 150mM NaCl/ 0.5% v/v IGEPAL/ 0.25% v/v Triton-X/ 10% v/v Glycerol) supplemented with 1× protease inhibitor cocktail (PIC-Roche, 04 693 116 001). After centrifugation, the cells were resuspended in 4ml Wash Buffer (WB: 10mM Tris-HCl pH 8/ 200mM NaCl) supplemented with 0.5mM DTT and incubated for 10min on a rotating wheel at 4$^{o}$ C. Nuclei were pelleted for 5min at 3000 rpm at 4$^{o}$ C and resuspended in 1.5ml RIPA buffer (10mM Tris-HCl pH 8/ 140mM NaCl/ 1% Triton-X/ 0.1% Sodium-Deoxycholate/ 0.1% SDS) freshly supplemented with 1× protease inhibitor cocktail and 1mM $CaCl_2$. Samples were equally split in 3 (1.5 ml tubes) and incubated for 10min in a water bath at 37$^{o}$C. MNase (Thermo Scientific EN0181) was diluted at 80U/µl in RIPA buffer freshly supplemented with 1× protease inhibitor cocktail and 1mM $CaCl_2$. The digestion was carried out by adding MNase at a final concentration of 6U per $10^6$ cells and incubation for 5min at 37$^{o}$C with occasional mixing. Digestions were stopped by placing tubes on ice and by adding 2×STOP Buffer (2% Triton/ 0.2% SDS/ 300mM NaCl/ 10mM EDTA). Digested chromatin was recovered by overnight rotation at 4$^{o}$C followed by centrifugation at 4$^{o}$ C at maximum speed for 10min. The chromatin was equally distributed in 1.5ml tubes and the crosslinking reversed overnight at 65°C in the presence of 1% SDS and shaking (800 rpm). The next day the samples were incubated for 2h at 56°C with 5µl Proteinase K (GEXPRK01B5 Eurobio) and processed through phenol/chlorophorm extraction and ethanol precipitation. DNA was resuspended in a total volume of 0.4ml UltraPure Nuclease free water. RNase digestion was carried out with 5µl RNase A (EN0531 Thermo Scientific) for 2h at 37$^{o}$C. The DNA was purified and precipitated again and resuspended in 51µl UltraPure Nuclease free water. 1µl of DNA was further diluted and the fragment sizes were analysed with a D1000 HS Screentape (Agilent 5067-5584) on an Agilent Tapestation.

*CLICK reaction and Streptavidin Pull-Down*: each DNA sample (50µl) was supplemented, in order, with the following buffers: 342.5µl of 100mM Potassium Phosphate Buffer (15433339

Fisher Scientific); 50µl of 10mM Biotin-TEG Azide (762024 Sigma) resuspended in DMSO; 7.5µl of pre-mixed Cupric Sulfate Pentahydrate (2.5µl at 20mM; C8027 Sigma) and THTPA (5µl at 50mM; 762342 Sigma); 25µl of 100mM Aminoguanidine Hydrochloride (396494 Sigma); 25µl of 100mM Sodium Ascorbate (A4034 Sigma). After briefly vortexing, the reactions were performed for 1.5h at room temperature protected from light, and DNA was subsequently ethanol-precipitated and resuspended in 400µl UltraPure water. 20µl were kept as input and the rest (380µl) were used for streptavidin pull down. For this, 25µl M-280 streptavidin magnetic beads (10292593 Thermo Scientific) per reaction were washed 3 times (10min rotation at room temperature per wash) with 1ml 1× Wash and Binding buffer (W&B); (2×W&B: 10mM Tris-HCl pH 7.5; 1mM EDTA; 2M NaCl; 0.2% Tween 20) and resuspended in 25µl 1× W&B buffer. EdU-labelled DNA was then pulled down with 380µl 2× W&B buffer and 25µl washed beads (1h at room temperature on a rotating wheel). Beads were separated by placing the tubes on a magnetic rack and the unbound fraction was stored at -20$^{o}$C. The beads were washed 3 times with 1ml W&B buffer (10min at room temperature on a rotating wheel). Washed beads were resuspended in 100µl TE/1% SDS and DNA was eluted with 20µl Proteinase K (1h at 65$^{o}$C with occasional vortexing). After adding TE to a final volume of 200µl, the DNA was phenol/chloroform-extracted, ethanol-precipitated and resuspended in 20µl UltraPure water.

*Library preparation and sequencing*: we used 10ng of DNA from the input samples and the whole pulled-down samples to prepare libraries as previously described (*1*). Sequencing (PE150) was performed by Novogene Co. Ltd. Only samples from experiments with non-detectable levels of DNA in control samples where biotin was replaced by DMSO during the CLICK reaction were used for library preparation.

**ChIP-seq**

For interphase, $2.5x10^6$ fixed cells (1% formaldehyde for 10min followed by 5min with 0.125 M Glycine) were resuspended in 2ml swelling buffer (25mM Hepes pH 7.95, 10mM KCl, 10mM EDTA; freshly supplemented with 1× protease inhibitor cocktail – 04 693 116 001 PIC-Roche, and 0.5% IGEPAL), incubated for 30min on ice, and passed 40 times in a dounce homogenizer to recover nuclei. For mitotic cells, the homogenization step was omitted. Mitotic and interphase cells were then treated in parallel to sonicate the chromatin in 300µl of TSE150 (0.1% SDS, 1% Triton X-100, 2mM EDTA, 20mM Tris-HCl pH8, 150mM NaCl; freshly supplemented with 1× PIC) using 1.5ml tubes (Diagenode) and a Bioruptor Pico (Diagenode; 7 cycles divided into 30 s ON–30 s OFF sub-cycles at maximum power, in circulating ice-cold water). After centrifugation (30min, full speed, 4 °C), the supernatant was pre-cleared for 2h with 50µl of protein G Sepharose beads (P3296-5 ML Sigma) 50% slurry, previously blocked with BSA (500 µg/ml; 5931665103 Roche) and yeast tRNA (1 µg/ml; AM7119 Invitrogen). 20µl were set apart (input) before over-night immunoprecipitation at 4°C on a rotating wheel with 4µl of anti-CTCF (Active Motif 61311) or 4µl of anti-SMC1 (Bethyl Laboratories A300-055A) antibodies in 500µl of TSE150. Protein G beads (25µL 50% slurry) were added for 4 h rotating on-wheel at 4 °C. Beads were pelleted and washed for 15min rotating on-wheel at 4 °C with 1ml of the following buffers: 3 washes with TSE150, 1 wash with TSE500 (as TSE150 but 500 mM NaCl), 1 wash with Washing buffer (10mM Tris-HCl pH8, 0.25M LiCl, 0.5% NP-40, 0.5% Na-deoxycholate, 1mM EDTA), and 2 washes with TE (10mM Tris-HCl pH8, 1mM EDTA). Elution was performed in 100µl of elution buffer (1% SDS, 10mM EDTA, 50mM Tris-HCl pH 8) for 15min at 65 °C after vigorous vortexing. Eluates were collected after centrifugation and beads rinsed in 150µl of TE-SDS1%. After centrifugation, the supernatant was pooled with the corresponding first eluate. For both immunoprecipitated and input chromatin, the crosslinking was reversed overnight at 65 °C, followed by proteinase K treatment, phenol/chloroform extraction and ethanol precipitation.

H3 ChIP-seq was performed on MNase-digested chromatin using $2.5x10^6$ cells fixed with formaldehyde as described above. Fixed cells were resuspended in 500µl of MNase buffer (50 mM Tris-HCl pH8, 1 mM CaCl2, 0.2% Triton X-100) supplemented with protease inhibitor cocktail (PIC; 04 693 116 001 Roche). Cells were pre-incubated for 10min at 37°C with 16U of MNase (Expressed in KUntiz, 1 Kunitz is equivalent to 10 gel units, NEB M0247S) added to the reaction. Cells were incubated for further 10min at 37°C, inverting the tubes occasionally. The reaction was stopped on ice by adding 500µl of 2×STOP buffer (2% Triton X-100, 0.2% SDS, 300mM NaCl, 10mM EDTA). Tubes were left rotating overnight on a wheel to allow diffusion of the digested fragments. The cell suspension was spun down and the supernatant stored at -80°C. Subsequently, 50µl of chromatin were brought to a final volume of 500µl with TSE150 (0.1% SDS, 1% Triton X-100, 2mM EDTA, 20mM Tris-HCl pH8, 150mM NaCl), freshly supplemented with protease inhibitor cocktail (PIC-Roche, 04 693 116 001), and ChIP was performed as described above using 5µg of anti-Histone H3 rabbit polyclonal (Abcam; ab1791).

All libraries were generated as previously described (*1*) and sequenced in the BioMics facility of the Institut Pasteur.

**Western Blot**

For Western Blot analysis cell pellets corresponding to $10^6$ cells were resuspended in 100µl Laemmli Sample Buffer (161-0737 BIO-RAD) containing β-mercaptoethanol. Lysed pellets were boiled for 10min at 95$^o$C and centrifuged for 10min at maximum speed at room temperature. Typically, 10µl per sample were loaded on 4-15% Mini-PROTEAN TGX Stain-

Free Gels (4568086 BIO-RAD) and run in 1×SDS-Running Buffer (250mM Tris/ 1.92M Glycine/ 1% SDS) at 10-20mA using the Mini-PROTEAN Tetra System (BIO-RAD). Proteins were transferred on nitrocellulose membranes (Amersham Protran 10600003) for 1h (or 3h for CTCF) at 300mA using the wet transfer system (BIO-RAD) in 1× Transfer Buffer (10× 0.25M Tris/ 1.92M Glycine) prepared with a final concentration of 20% Ethanol. Membranes were blocked in PBST (PBS 0.1% Tween-20) 5% BSA for 1h at room temperature and incubated over night at 4 $^{o}$C with primary antibodies diluted in PBST 5% BSA (1:5000 anti-H3 Abcam ab1791; 1:500 anti-CTCF Active Motif 61311). Excess antibodies were washed off with PBST (5 washes, 5 min each) and incubated for 1h at room temperature in secondary antibodies HRP-conjugated diluted in PBST 5% BSA (1:10000 anti-Rabbit IgG-HRP Thermo Fisher RB230254). Membranes were washed 5 times, 10min each at room temperature and incubated with PIERCE ECL2 Western Blotting Substrate (80196 Thermo Scientific) 5min in dark. After excess reagent was removed, proteins were visualised using the BIO-RAD Chemidoc MP Imaging System and processed using the Image Lab Software (BIO-RAD).

**Imaging**

Immunostaining of ES, C2C12 and NIH3T3 cells: cells were plated at a density of $2\times10^4$ on IBIDI hitreat plates coated overnight with poly-L-ornithine (0.01%; Sigma, P4957) at 4 °C, washed and coated 2h with laminin (10μg/ml in PBS; Millipore, CC095). Two days post-seeding, cells were washed twice with PBS, fixed for 10 min at room temperature with PBS 4% Formaldehyde (Thermo, 28908), washed twice in PBS and permeabilised with PBS/0.1% v/v Triton X-100 supplemented with 3% of donkey serum (Sigma, D9663) for 15min at room temperature. Incubation with primary anti-CTCF (1:500; Active Motif 61311) and anti-SMC1 (1:500; Bethyl Laboratories A300-055A) antibodies were performed in PBS with 3% donkey serum. After three washes in PBS/0.1% Triton X-100, Alexa Fluor 594 AffiniPure Donkey Anti-Rabbit secondary antibodies (Jackson ImmunoResearch, 711-585-152) were were applied for 2h at room temperature (2µg/ml). Cells were washed three times in PBS/0.1% v/v Triton X-100, nuclei counterstained with 4',6-diamidino-2- phenylindole (DAPI; Sigma, D9542), and imaged with a LSM800 Zeiss microscope using a 64× or 40× oil immersion objective.

*Live imaging*: CTCF-aid ES cells also carry a GFP fused in frame with CTCF-Aid (*6*), enabling live imaging of CTCF. To do this, the cells were cultured on IBIDI plates in phenol red-free medium (Gibco, 31053-028) and incubated with 500nM Hoechst-33342 for 20min before imaging, which was performed at 37 °C in a humidified atmosphere (7% $CO_2$). Images were acquired with a 63× oil immersion objective on a Zeiss AxioObserver Z1 microscope equipped with a Yokogawa CSUX1 spinning-disk confocal scanner, a Hamamatsu EMCCD ImageEM X2 camera using the Volocity acquisition software.

*Immunostaining of embryos*: All experiments were conducted according to the French and European regulations on care and protection of laboratory animals (EC Directive 86/609, French Law 2001-486 issued on June 6, 2001) and were approved by the Institut Pasteur ethics committee (n° dha180008). CD1 (Charles River Laboratories, France) embryos were recovered at E3.5 (mid-blastocyst stage) and cultured for 6h in EmbryoMax KSOM (Millipore Bioscience, MR-121D) with or without 5µM Nocadazol (Tocris Biotechne, 1228) to enrich for mitotic cells. Embryos were fixed for 15min at room temperature in 4% paraformaldehyde (Euromedex, 15714), rinsed with 1×PBS and incubated in 1×PBS, 0.1% Triton-X100, 10% donkey serum for 30min at RT. Embryos were then incubated with primary antibodies overnight at 4°C and with secondary antibodies for 1h at RT in 1×PBS, 0.1% triton-X100, 10%

donkey serum. The following antibodies were used: anti-Nanog (1:200; eBioscience 14-5761-80), anti-CTCF (1:300; Active Motif 61311), donkey anti-rat Alexa Fluor 488 (1:300; Invitrogen A21208) and donkey anti-rabbit Alexa Fluor 546 (1:300; Invitrogen A10040). Nuclei were counterstained with Hoechst 33342 (1.6µM; Sigma Aldrich 14533), and embryos were placed individually in glass bottom microwell dishes (MatTek Corporation P35G-1.5-20-C) in 1×PBS. Fluorescent images were obtained using a confocal laser-scanning microscope (LSM800; Zeiss) with the objective Plan-apochromat 20× /NA 0.8, speed 6, pinhole 1 airy unit, and laser intensities suited for optical section thickness of 2µm. Images were analyzed and processed using Fiji and Photoshop CS6 softwares. Pictures correspond to a projection of 5 confocal optical slices.

**Computational Methods**

**Data and availability**

Samples are summarized in Table S1. Briefly for ChIP-seq, we sequenced 3 x Interphase and 3 x Mitosis wild-type CTCF; 2 x Interphase and 2 x Mitosis wild-type SMC1; 2 x Interphase and 2 x Mitosis CTCF in CTCF-aid for -IAA and +IAA (2h); 1 x interphase and 1 x mitosis of CTCF in C2C12 and NIH3T3 cells. For MINCE-seq we sequenced 2 x input and IP pairs for pulse and chase samples. Finally MNase H3 ChIP-seq in CTCF-aid 2 x Interphase and 2 x mitosis for both -IAA and +IAA (2h). We previously generated wild-type MNase-seq: 3 x Interphase and 3 x Mitosis; MNase H3 ChIP-seq: 2 x Interphase and 2 x Mitotsis; and ATAC-seq: 2 x Interphase and 2 x Mitosis. All previously generated datasets are described (1) and are available from GEO accession: GSE122589.

**CTCF and SMC1 binding in interphase and in mitosis.**

*Data Processing:* reads were aligned with Bowtie 2 (7) to the mm10 genome, with options "-k 10". Reads were additionally filtered for those with a single discovered alignment (in Bowtie 2 "k" mode this is mapping quality = 255) and an edit distance less than 4. All libraries were constructed with custom unique molecular identifier barcode as described before (1). Therefore, reads aligning with identical position, strand and barcode were treated as duplicates and collapsed.

*Peak calling:* we used a strategy previously described (1), where peaks were called against relevant inputs/controls for all samples using MACS2 (8) with "callpeak -q 0.2 -g mm". Peaks intersecting with the mm10 blacklist (9) were excluded along with those on chrM and chrY. To determine a set of candidate binding regions for each factor we merged all peaks in interphase and mitosis. Peaks were further filtered to have MACS2 FDR < 0.01 in both replicates in either interphase or mitosis, and a height > 0.8 reads per million (RPM) in at least one sample (a height of ~16 raw reads at our mean mapped read depth of ~20 million reads).

*Bookmarking analysis:* we used a strategy previously described (1), where we take an approach similar to differential expression analysis in RNA-seq to determine the set of peaks bookmarked CTCF and SMC1. We combine the read counts for input and ChIP samples for a given TF in a single generalized linear model (GLM) aimed at assessing the difference between input and ChIP signal in both interphase and mitosis at each peak. We implement this GLM using DESeq2 (10). We expect that nearly all peaks will have significant differences between input and ChIP, and that the majority of peaks will have significantly different occupancy between interphase and mitosis. Therefore, we set the size factors to the total mapped reads of each sample. We encode ChIP versus Input by a factor ChipTF and mitosis samples by a factor ChipM, and then we ran Wald tests on the model ~ChipTF + ChipM. At significance FDR < 0.05 and without independent filtering, we tested ChipTF (the fold change of interphase over input), the sum ChipTF + ChipM (the fold change of mitosis over input), and finally ChipM the difference between interphase and mitosis. To determine the set of bookmarked peaks we required that a peak had differential occupancy (ChipTF + ChipM, FDR < 0.05), that both mitosis replicates had MACS2 FDR < 0.01 with one mitosis replicate had MACS2 FDR < $10^{-10}$. Combined, this is a conservative strategy that helps ameliorate the effects of contamination from interphase cells (1). The significance of ChipM was further used to classify three types of bookmarking peaks with either higher enrichment in interphase (BI), in mitosis (BM), or similar in both phases (BS). This resulted in:

CTCF: Lost (L): 22,550, BI: 18,723, BS: 10,456, BM: 360

SMC1: L: 17,251

*Visualization and normalization of ChIP-seq data*: single end read data was extended to mean fragment length (~200bp for CTCF-aid samples and ~120bp otherwise). For CTCF, the single base pair at the center of the inferred fragment was marked and normalized to units of reads per million (RPM), with the exception of CTCF-aid mitosis, interphase and mitosis comparison between E14Tg2a, C2C12 and NIH3T3 cells, due to low signal in some of these samples. In these cases, the entire fragment was marked and normalized to be comparable to single base heatmaps. SMC1 heatmaps were generated by marking entire fragments. All heatmaps are ordered by descending peak height and are at resolution of 100 sites per pixel. Heatmaps color scale is linear (inferno colormap) scaled to 0.2 of wild-type interphase (or mitosis for Fig. S4G) peak maximum.

*Transcription Factor Motif Detection*: to discover CTCF binding motifs we scanned the mm10 genome using FIMO (11) for motif MA0139.1 with parameters “--thresh 1e-3 --max-stored-scores 50000000” and supplied a 0-order Markov background file describing the relative nucleotide frequencies in the mm10 assembly. Motifs were intersected with peaks. Metaplots, V-plots and heatmaps are centered on the best scoring motif within each peak.

**Nucleosome Positioning MNase-seq, MNase-H3-ChIP-seq, MINCE-seq.**

*Data Processing:* paired end reads were trimmed by aligning read pairs to discover regions of reverse complementarity surrounded by our custom adapters. Alignment and trimming were performed with the BioSequences package for Julia 0.6 (12). Reads were aligned to mm9 genome using Bowtie 2 (7) with options “-k 10 -I 0 -X 1000 --no-discordant --no-mixed”, and filtered for reads with a single alignment mean edit distance less than 4 between read pairs. For all libraries with barcoded adapters, identical barcodes with identical position were collapsed.

*Bias correction*: in all MNase samples we estimated and corrected the MNase cutting bias. We use our previously applied approach (1), which is a simplified version of seqOutBias (13). We evaluated the relative rates of occurrence of the k-mer at the end of each read and the total occurrence of that k-mer in the genome. Specifically, we took the two 6-mers lying over the end coordinates of each read, such that each 6-mer was composed of two 3-mers, one lying within the read and one lying outside. We counted total k-mers in the genome using the BioSequences package in Julia 0.6 (12). To assess an appropriate correction, we averaged the k-mers at the end of each read in a position weight matrix (PWM), we found the left

PWM to be approximately equal to the reverse complement of the right PWM, and so we calculated a correction based on the left k-mer and the reverse complement of the right k-mer. If $\gamma_i$ is the rate of occurrence of k-mer $k_i$ in the genome, $\rho_i$ is the rate of occurrence of the same k-mer at MNase cutsites, and $k_L$ is the left k-mer and $k_{R^\dagger}$ is the reverse complement of the right k-mer, then in all analysis each fragment is weighted by: $\sqrt{\rho_L \rho_{R^\dagger} / \gamma_L \gamma_{R^\dagger}}$. We employed a pseudocount of 100 in calculating k-mer rates.

*MNase fragment size selection*: we employed V-plots (Figs. 1A, 2C, 4C and S4A) over features to conservatively and robustly select fragment windows for footprinting and evaluating nucleosome signal. For MNase samples, we selected fragments less than 100bp for footprints and fragments in the interval [140, 200] bp for nucleosomes. For MINCE-seq libraries which exhibited a different fragment size distribution we used fragments in the interval [120, 200] bp for nucleosomes.

*Nucleosome normalization*: for wild-type analysis we calculated the rate of mid-points of nucleosomal size fragments per billion (mid-points per billion - MPB) at each base pair per site for heatmaps and single loci, and averaged over all sites for metaplots and V-plots. For MINCE-seq and CTCF-aid MNase H3 ChIP-seq which exhibited more variation in fragment size distributions we calculated the MPB relative to total nucleosomal size fragments. To focus on differences of nucleosomal organization in MINCE-seq, we normalized total nucleosomal fragments within [-1kb, +1kb] of the center of each feature in pulse and chase to be equal to the relevant input. Similarly, for CTCF-aid samples we normalized total nucleosomes per feature in +IAA to be equal to -IAA, and in (Fig. 4D, Fig. S4H) we further normalised to total wild-type interphase nucleosomes.

*Nucleosome visualization:* all heatmaps mark the midpoint of nucleosomal sizes MNase or MNase-H3 nucleosomal fragments in MPB, displayed at 100-site per pixel resolution with inferno colormap scaled to wild-type interphase MNase or MNase H3 maximum as appropriate. Metaplots are calculated at base pair resolution by the mean MPB per base pair per site surrounding each feature (visualized as the point clouds in Figs. 2D, 3A and 4D); we then apply Gaussian process regression to smooth and model nucleosome positions as described below.

*Nucleosome positioning regression:* to assess the statistics of the mean nucleosome signal over a set of regions we employed Gaussian process regression (15) on MNase and MNase H3 metaplots. We used a squared exponential covariance function and selected hyper-parameters for signal variance, length scale and noise variance, optimized on [-500, 500] bp interval surrounding the central point of each feature. For all but CTCF-aid data, this contains the signal of primary significance and importantly the covariances are relatively stationary over this region as compared to outside where length and noise scales change as the data loses coherence. For CTCF-aid +IAA interphase and +/- IAA mitosis data the nucleosomes roll inwards over the region of maximum MNase-cutting bias (visible as V in V-plot Fig. 2C), causing an artificial break in the +/-1 nucleosomes with altered covariance. Therefore we optimized hyperparameters on the union [-500, -100] U [100, 500] bp. We then use these optimized hyperparameters to predict over the full region. We selected hyperparameters by sparse Gaussian process regression employing a variational approximation to the marginal likelihood (14; https://github.com/SheffieldML/GPy), with an initial inducing inputs every 10bp. To account for the overdispersed count-data nature of the nucleosome fragment midpoints, we employ a log transform to stabilize variance and then assume a Gaussian likelihood for Gaussian process regression. We transform the total midpoints $m_i$ at site $i$ data by $y_i = \log(\alpha m_i + \beta)$, we select $\alpha = 10^3$and $\beta = 1$. We make predictions and invert the transform, $\hat{m}_i = (e^{\hat{y}_i} - \beta)/\alpha$ where $\hat{y}_i$ is the Gaussian process mean, thus $\hat{m}_i$ reflects the median of the transformed random variable and we plot this median.

*Nucleosome positioning*: to assess the position of the +/- 1 nucleosomes and their dependence on CTCF occupancy, we took the nucleosome fragment midpoints in [-230, -70] bp for the -1 nucleosome and [70, 230] bp relative to CTCF motif center in 100 site bins descending with CTCF peak size. For each bin we evaluated the median of the empirical cumulative density function within each region. If $m_{ij}$ is the total number of nucleosome fragment midpoints at base $i$ in site $j$, we calculate $\hat{F}_j(i) = \left(\sum_{k \leq i} m_{kj}\right)/\left(\sum_k m_{kj}\right)$, and take the median as the smallest $i$ such that $\hat{F}_j(i) \geq 0.5$. We then smooth the medians per 100-site bins by Gaussian process regression. To asses nucleosome positioning over a NOA averaged over a set of sites we call the maxima of the Gaussian process median described above.

*Nucleosome spectral density*: to assess nucleosome periodicity we report the spectral density (Fig. 3B) of the covariance function with optimized hyperparameters. For the squared exponential kernel, frequency $s$ and lengthscale $\ell$ this is given by $S(s, \ell) = \sqrt{2\pi\ell^2}\exp(2\pi^2 s^2 \ell^2)$ (15), we evaluate spectral density at period p = 180, s = 1/180 which corresponds to nucleosome plus linker.

**Chromatin accessibility in interphase and in mitosis.**

Paired end reads were trimmed by aligning read pairs to discover regions of reverse complementarity surrounded by Nextera sequencing adapters for ATAC-seq. Alignment and trimming was performed with the BioSequences package for Julia 0.6 (12). Reads were aligned to mm10 genome using Bowtie 2 (7) with options "-k 10 -I 0 -X 1000 --no-discordant --no-mixed", and filtered for reads with a single alignment mean edit distance less than 4 between read pairs. To generate heatmaps, the two end points (cut sites) of fragments in the 0-100 bp range, shifted inward by +/- 4bp as recommended (16) and piled at base pair resolution. Heatmaps are visualized at a resolution of 100 sites per pixel using inferno colormap scaled to 0.5 maximum interphase signal.

**Genomic features.**

All heatmaps are metaplots are centered on CTCF maximal motifs (Table S2); SMC1 summits (Table S3); Esrrb motifs (Fig. 3A) for Esrrb bookmarked regions were Esrrb dictates nucleosomal organization as previously defined (1); Oct4/Sox2 composite motifs (Fig. 3A) at Oct4 and Sox2 interphase binding regions as defined (1); P300 binding regions centered of summits derived from the Encode portal (9, 17), experiment identifier: ENCSR000CCD, file: ENCFF179FJG. We further restricted P300 binding regions by a set of ChromHMM (18) defined as ES enhancers (19; https://github.com/guifengwei/ChromHMM_mESC_mm1). We retain P300 binding regions that intersected as regions annotated as "Enhancer", "Strong Enhancer" or "Weak/Poised Enhancer" and no other ChromHMM categories. To asses CTCF binding at different chromatin states (Fig. 2D) we used the same ChromHMM data and selected insulators, the enhancer set described above, active promoters and active gene bodies ("Transcription Elongation").
